# Glycocalyx-Directed Enzymatic Hydrogels Unlocks Fibroblast Regeneration to Promote Diabetic Wound Healing

**DOI:** 10.64898/2026.07.23.736655

**Authors:** Vignesh Ganesan, Iffat Jahan, Anisha Karmakar, Swapnil Raut, Sarbajeet Dutta, Md. Asrafuddoza Hazari, Jayashri Pandya, Renuka Munshi, Dipti Kumbhar, Lokesh Kumar Bhatt, Shamik Sen

## Abstract

Chronic diabetic wounds remain a major clinical challenge because current therapies address infection or supplement growth factors without correcting the cellular dysfunction that prevents regeneration. We identify pathological glycocalyx thickening in diabetic dermal fibroblasts (DDFs) as a driver of elevated caspase 3, 8, and 9 expression and heightened apoptosis — deficits that stall wound repair. Cleavage of sialic acid residues by neuraminidase (NMase) reverses this dysfunction, restoring fibroblast migration, proliferation, and contractility. We engineer a photo-crosslinked hybrid hydrogel combining methacrylated gelatin (GelMA) with high-molecular-weight methacrylated chitosan (HMW ChMA). ChMA increases storage modulus 8–12-fold, reduces pore size, and confers antibacterial activity against gram-positive and gram-negative bacteria, addressing infection susceptibility. The fortified HMW hybrid (HMWH) network enables sustained, localized NMase delivery that outperforms GelMA alone in resisting degradation and controlling release kinetics. NMase-loaded HMWH (N-HMWH) gels enhance DDF proliferation and migration in vitro, correlating with reduced focal adhesion size and increased turnover. In a diabetic rat model, N-HMWH patches achieve superior wound closure, outperforming EGF therapy, with robust epidermal regeneration, neovascularization, and collagen deposition. This work establishes glycocalyx-targeting hydrogels as a new class of wound therapeutics addressing the root cause of diabetic fibroblast failure, not just compensating with growth factors.

## 1. Introduction

Diabetic wounds remain a major clinical challenge due to multiple factors such as poor vascularization, prolonged inflammation, and cellular dysfunction that collectively impair tissue regeneration. Under normal physiological conditions, wound healing is initiated by tissue injury and proceeds through a series of phases starting with clotting, inflammation, proliferative, and remodeling phases, driven by coordinated cytokine and growth factor signaling [1] [2] [3]. In diabetes, however, this process is profoundly dysregulated, resulting in delayed cell recruitment, reduced matrix deposition, and increased susceptibility to infection [4] [5]. These problems root from various impediments such as impaired perfusion, elevated oxidative stress, and increased apoptosis which frequently lead to the development of chronic, non-healing ulcers [6] [7].

Fibroblasts plays an integral part in wound repair by migrating into the wound bed, synthesizing extracellular matrix (ECM), and generating contractile forces necessary for wound closure [8] [9] [10] [11]Diabetic fibroblasts possess lower activity, including reduced migratory capacity, impaired collagen synthesis, and diminished contractility, which compromises their ability to form granulation tissue and mediate matrix remodeling [12] [13] [14]. Current therapeutic approaches for diabetic wound care largely focus on infection control through antibacterial or antifungal formulations, or on the exogenous administration of growth factors such as fibroblast growth factor (FGF), epidermal growth factor (EGF), and vascular endothelial growth factor (VEGF) [15] [16] [17] [18].

Recent studies have identified the cellular glycocalyx as a key regulator of cell–matrix interactions, mechanotransduction, and migratory behavior [19] [20] [21] [22]. The glycocalyx is a dense layer composed of proteoglycans, glycoproteins, and glycosaminoglycans that modulates force transmission between the extracellular environment and the cytoskeleton [19] [23] [24] [25]. In diabetic fibroblasts, an abnormally thickened glycocalyx has been reported, which correlates with impaired focal adhesion dynamics, attenuated mechanosensing, and reduced cellular contractility [26]. Enzymatic reduction of glycocalyx thickness using neuraminidase, which cleaves terminal sialic acid residues, has been shown to restore fibroblast migration, enhance collagen deposition, and increase contractile force generation. These findings suggest that targeted glycocalyx modulation represents a promising strategy for restoring fibroblast function in diabetic wound healing contractility [26].

In this study, we have engineered a bioactive hybrid hydrogel composed of gelatin and chitosan as a wound dressing and carrier for neuraminidase. Gelatin and chitosan were used due to their inherent biocompatibility, biodegradability, and established utility in wound healing applications. To improve the mechanical stability and tunability of the hydrogel network, both polymers were chemically modified via methacrylation, enabling covalent crosslinking and controlled modulation of network density. The resulting methacrylated gelatin/chitosan hybrid hydrogel provides a viable environment for cell to migrate and divide while facilitating NMase release at the wound site. The therapeutic potential of neuraminidase-loaded hydrogels was demonstrated both in vitro using patient derived fibroblasts and in vivo in diabetic wound models. Neuraminidase-loaded hydrogels led to enhanced migration and proliferation compared to cells on control hydrogels, indicating restoration of key functional behaviors essential for wound repair. In vivo, treatment with neuraminidase-loaded hydrogels resulted in accelerated wound closure, accompanied by enhanced collagen deposition, neovascularization and re-epithelialization. Overall, this work demonstrates that enzymatic modulation of the glycocalyx, delivered through a hydrogel system can improve fibroblast function and promote tissue regeneration in diabetic wounds. By directly targeting the underlying cause that impairs fibroblast activity during healing, this approach offers a distinct alternative to conventional growth factor-based therapies and sheds light on more alternative therapies such as the potential of glycocalyx focused strategies in advanced wound healing biomaterials.

## 2. Materials and Methods

### 2.1 Isolation and Culture of Human dermal fibroblasts

In-vitro experiments were performed using primary human dermal fibroblasts isolated from human skin samples collected from the BYL Nair hospital, Mumbai, India. Only discarded skin samples were collected after the consent of diabetic and non-diabetic patients (10 diabetics, 10 non-diabetics, age: 40-60) undergoing any elective surgery. The skin samples were transported in plain DMEM media supplemented with 1% Penstrep (Gibco, Cat# 15140122) and 1% Funigizone (Invitrogen, Cat#15290018) at 4°C; fibroblasts were isolated within 3-4 hours from the time of isolation. Ethical clearance for using the tissues was approved by the ethical committees from both the institutes. Fibroblasts were isolated as per previously established protocols [26] [27] [28] [29]. After removing fat present in the tissue samples using surgical scissors, tissues were cut into smaller pieces 1-2 cm in size. The samples were then incubated with dispase (Himedia, Cat#42613-33-2) solution of 2.5 U/ml at 4° C overnight to separate the epidermis from the dermis. The dermis was further sliced into tiny pieces ∼1 mm size using a scalpel and then incubated in collagenase (Himedia, Cat# 17104019) solution in 60 mm Petri dishes at a concentration of 1600U/ml for 150 minutes at 37° C in a CO_2_ incubator; the petri dishes were taken out and gently shaken every 30 minutes. Next, a 70-micron cell strainer was used to separate the cells from the collagenase treated tissue samples. The filtered solution containing the cells was centrifuged at 1500rpm for 5 minutes. The pellet was then used for seeding cells in 100 mm dishes, expanded, and stored in frozen vials with passage number 1-3. The cells were cultured in DMEM high glucose media (Gibco, Cat#12800-017) supplemented with 10% FBS, (Gibco, South American origin, Cat # 10270-106), 1% pen strep,1% Sodium Pyruvate (Gibco, Cat#11360070) and 1mM Glutamine (Gibco, Cat# 35050061). The cells were passaged when they are 70-80 % confluent using 0.05% trypsin and 0.02% ETDA (Himedia, Cat#TCL140), and were used for experiments for a passage number of 2-6.

### 2.2 Preparation and characterization of methacrylated gelatin and methacrylated chitosan

Methacrylation of gelatin (porcine, Sigma, Cat#G2526) was done as described elsewhere [30] [31] [32] [33] [34], by first dissolving the gelatin in PBS to make a 10 % solution at 55°C. After the gelatin was homogeneously dissolved in PBS, the temperature was brought down to 53°C and 10ml of Methacrylic Anhydride (MA) was added dropwise to the solution under constant stirring. The experiment was performed inside a fume hood under dark conditions since MA is toxic and to prevent any crosslinking. The reaction was continued for 2 hours and 4-fold volume of PBS was added in the end to arrest the reaction. The final solution was transferred to a 14kDa dialysis bag and dialysis done for 7 days using distilled water at 40°C. The solution after dialysis was freeze-dried to form a sponge of methacrylated gelatin (GelMA). Similarly, the methacrylation of chitosan was done by adding deacetylated chitosan in 2% (v/v) acetic acid solution to make a 3% (wt/v) chitosan solution and placed in a shaker overnight to completely dissolve the chitosan. Two types of chitosan were used in making the gels having different molecular weight, a low molecular weight (LMW) chitosan (50-190 kDa, Sigma Cat#448869-50G) and a high molecular weight (HMW) chitosan (190-330 kDa, Sigma Cat#448877-50G). Deacetylated chitosan was prepared by stirring the chitosan in 1M NaOH solution for 4 hours at 60°C. The chitosan was then completely dried at 50°C and dissolved in 1M acetic acid solution, with the pH of the solution adjusted to 7 by gradually adding 5N NaOH. The chitosan solution having a pH of 7 was centrifuged at 4000 rpm, with the precipitated chitosan was collected and lyophilized. This chitosan was then dissolved in 1 M acetic acid solution and methacrylation was done by adding 340 mg of methacrylic anhydride to the solution dropwise under rigorous stirring at RT. After 4 hours, the solution was transferred to a dialysis bag and dialysed for 5 days and later lyophilized. The lyophilized methacrylated chitosan (LMW/HMW ChMA) was stored at -20 C. The degree of methacrylation (DM) of the GelMA and ChMA was determined using proton NMR., where GelMA was dissolved in deuterium oxide and ChMA was dissolved in deuterium chloride/deuterium oxide solution. For GelMA, the DM was calculated by measuring the ratio of the area under the curve for the peaks at 2.8-3.0 ppm (represent lysine methylene) in GelMA spectrum to the peaks at 7.1-7.4 ppm (represent phenylalanine) in the gelatin spectrum [33]. Similarly, for ChMA it was the ratio of the peaks at 2.8-3.1 ppm (H2-H6 in benzene groups) to the peaks at 5.5-6.1 ppm (methylene peaks) [35]. The degree of deacetylation of chitosan was measured from peaks solid state FTIR from the ratio of the absorbance at wavenumber 1650 and 3450 cm^-1^ [35].

### 2.3 Preparation and characterization of GelMA and GelMA/ChMA hybrid gels

To prepare GelMA-ChMA hybrid gels, lyophilized ChMA (methacrylated chitosan) was dissolved in 1mM acetic acid solution to form a 1%(wt/v) solution using a shaker at 37°C for overnight. Once the ChMA was completely dissolved, GelMA was added such the final concentration of GelMA was 15% (w/v). Both LMW and HMW ChMA were used to fabricate the hybrid gels which we hereafter refer to as LMWH and HMWH gels respectively. The precursor gel solutions were mixed with Irgacure (2595) photo-initiator (15% w/v) 1:10 ratio of Irgacure and GelMA. The gels were polymerized by 365 nm UV exposure for 1 minute. Upon polymerization, gels were washed with PBS and UV sterilized for 30 minutes. The porosity of the hydrogels was measured from images taken using the field emission gun scanning electron microscopy. Small (5mm x 5mm x 2mm) blocks of hydrogels were mounted on stubs which were dipped in liquid nitrogen (-200 to -192 °C) and fractured. The samples were then sublimated at -85°C before coating with platinum for 30 second at 10mA. Images were taken at 10kV at a magnification of 10,000X and 20,000X. Physical properties of hydrogels were measured using a flat plate rheometer (Anton Paar) using disc shaped gels with radius of 1 cm and thickness of 4-5mm. A shear strain of 0.1 to 100 was applied to the gels at a frequency of 1

Hz to obtain the linear viscoelastic range. Swelling and degradation kinetics of the gels were studied by immersing the gels in 2µM collagenase solution and weighing them at regular time intervals. For assessing anti-bacterial activity, 5mm gel disks were prepared and incubated for 16 hours on E.Coli and S.Aureus bacteria spread agar gels and the zone of inhibition quantified. The release kinetics of neuraminidase (NMase) from the hydrogels was studied by measuring the cumulative release of the NMase. Similar to the degradation studies, the gels were loaded with NMase and submerged in 5 ml PBS. 2ml of the PBS was collected and replenished with fresh PBS at regular time intervals. To measure the amount of NMase present in the samples, we used MUNANA (2’-(4-Methylumbelliferyl)-α-D-N-acetylneuraminic acid) assay, a fluorescence-based assay where the NMase reacts with MUNANA to give a fluorescent product 4-MU (4-methylumbelliferone). To test the antibacterial property that chitosan inherently possesses, the disk diffusion assay was performed where 5 mm disk shaped hydrogel were made and placed on LB agar plate spread with bacteria. The plates were kept in the incubator for 16 hours with the temperature maintained at 37°C. Staphylococcus aureus MTC 96 and E.Coli K12 bacteria strains were used in the anti-bacterial studies with Ampicillin B and Kanamycin used as the positive control for these strains, respectively.

### 2.4 In-vitro studies on bioactive gels

Cells were seeded on 12 mm glass coverslips coated with the gels. All experiments were done 24 hours after seeding. For motility, time-lapse imaging was performed at 15-minute intervals for 12-hour duration. The analysis was done in ImageJ using cell tracking plugin. For immunostaining, the cells were fixed using 4% Paraformaldehyde (PFA) in PBS solution for 20 minutes. After fixing the cells were permeabilized using 0.5 % Triton-X solution for 5 minutes followed by 3 gentle washes of PBS. Blocking was done using 2% BSA solution for 1 hour. Following this, the primary antibody was added in the plates and incubated at 4°C overnight. The primary antibody was removed and the plates were washed 3 times for 10 minutes and then secondary antibody was added. Images were taken at 10X, 20X, and 63X.

### 2.5 Inducing diabetes in rats

Male Sprague Dawley rats (3-4-week-old, 150 ± 30g) were purchased from National Institute of Biosciences in Pune, India. Animals were housed in controlled temperature and humidity with a 12h light-dark cycle. All the experimental protocols were reviewed and approved by the Institutional Animal Ethics Committee (Approval Number: CCSEA/IAEC P-432023) and the experiments were carried out following the "Committee for Control and Supervision of Experiments on Animals" (CCSEA) guidelines. After one week of acclimatization and feeding a normal diet, animals were shifted to high-fat diet (60 kcal% fat). At the end of 4^th^ week, rats were fasted overnight and subsequently injected with freshly prepared streptozotocin (STZ, 30mg/kg, i.p.). The STZ solution was prepared using 0.1M citrate buffer (pH 4.5). Rats were continued on high-fat diet for the next 4 weeks. At 4^th^ week after STZ injection (8^th^ week of study), rats with blood glucose levels exceeding 220–250 mg/dL were classified as diabetic. Blood insulin concentration was measured using ELISA kit (Krishgen biosystems). Rats with blood glucose exceeding 220-250 mg/dl were randomly divided into six groups (n=6).

### 2.6 Wound healing Experiment

Following randomization, after 8 weeks, animals were anesthetized with ketamine and xylazine at a dose of 80mg/kg and 10mg/kg, respectively, for wound induction. Prior to the infliction of wounds, the dorsal fur of the animals was shaved, and the skin was disinfected using 70% ethanol. A standardized full-thickness, open excision wound of 1×1cm^2^ was made using scissors. Treatments were applied topically once immediately after making excisions, and wounds were wrapped with Tegaderm films. Lyophilized gel patches of the dimension of 1×1cm^2^ were rehydrated in PBS for 10 minutes before being applied to the wounds. The animals were divided in various treatment group as follows: Group 1 (Diabetic control, HFD + STZ + incision wound); Group 2 (EGF: HFD + STZ + incision wound + Regen-D (EGF)); Group 3 (GelMA: HFD + STZ + incision wound + GelMA gel); Group 4 (HMWH: HFD + STZ + incision wound + HMWH gel); Group 5 (N-GelMA: HFD + STZ + incision wound + N-GelMA gel); Group 6 (N-HMWH: HFD + STZ + incision wound + N-HMWH gel). In diabetic wounds, reduction in wound area over time predicts healing. On days 0, 3, 7, 10, and 14, the wound area of the experimental animals was photographed using digital camera and wound size was measured to evaluate the healing progress. The tegaderm films were removed after 3 days once the gels were strongly adhered to the wound surface. Wound assessments performed on Day 7 and Day 14 are critical indicators of healing process. In addition to wound area, the degree of wound closure and wound size were measured using Image J software. Percentage wound contraction was calculated as per the formula (% wound area): (Wo [original wound area]- Wt [wound area at specified time point]) / Wo [original wound area] × 100. When the area affected by the wound equaled the area at time point zero, the wound repair was said to be completed.

### 2.7 Histological analysis and ELISA

Animals were euthanized on Day 14 after wound induction. Skin at the wound sites was sampled at 5mm from the wound surface and was fixed in 10% neutral buffer formalin solution, followed by dehydration and paraffin embedding. Using microtome, the skin tissues were sectioned at 5µm thickness. The sections were then stained using Hematoxylin and Eosin and Masson’s trichome stain to study the extent of wound healing and tissue regeneration by evaluating epidermal thickness, neovascularization, and collagen formation. The slides of Masson’s trichome stain were photomicrographed using a digital camera attached to a light microscope (Motic Microscopes). ImageJ software was used to evaluate the percentage of blue staining, in the whole area of each image, with the dye indicating the presence of collagen fibres in the tissue. Microvessel count/high-power field was performed to evaluate neovascularization due to treatments. The concentration of plasma inflammatory cytokines (TNF-α and IL-6) levels were measured using commercially available ELISA kits (Krishgen biosystems) as per manufacturer’s protocol.

### 2.8 Statistical Analysis

All statistical analysis was done using the origin 2021 version. One-way ANOVA was performed for multiple groups and the means were compared using Tukey and Fisher tests. Within two group t-test was using the find the significance. Asterisks were used for the assigned p-values, * for p<=0.05, ** for p<=0.01 and *** for p<=0.001.

## 3. Results

### 3.1 Neuraminidase (NMase) treatment inhibits apoptosis and increases motility of diabetic dermal fibroblasts (DDFs)

Healthy dermal fibroblasts (HDFs) and diabetic dermal fibroblasts (DDFs) isolated from patient skin samples were used to perform in-vitro experiments to study their migratory and apoptotic behavior (Supp. Fig. 1A, B). In line with our previous findings [36], DDFs were more well spread and elongated than HDFs (Fig. 1A, B), with higher lectin intensity of DDFs indicative of a bulkier glycocalyx in these cells (Fig. 1B). To test the sensitivity of HDFs and DDFs to injury, a scratch wound assay was performed wherein after introducing the scratch, the dislodged cells were collected and seeded on poly-L-lysine-coated coverslips. Subsequently, after fixation, cells were stained for the apoptotic marker caspase 8 (Fig. 1C(i)). There was a substantial increase in the number of caspase 8 positive cells among DDFs (Fig. 1Cii, iii). Consistent with these observations, expression of caspase 3, 8 and 9 were elevated in DDFs compared to HDFs (Fig. 1Di). Treatment with caspase inhibitor Z-VAD-FMK (hereafter referred to as CI) led to substantial drop in expression of all the three genes (Fig. 1Dii).

**Figure 1.**
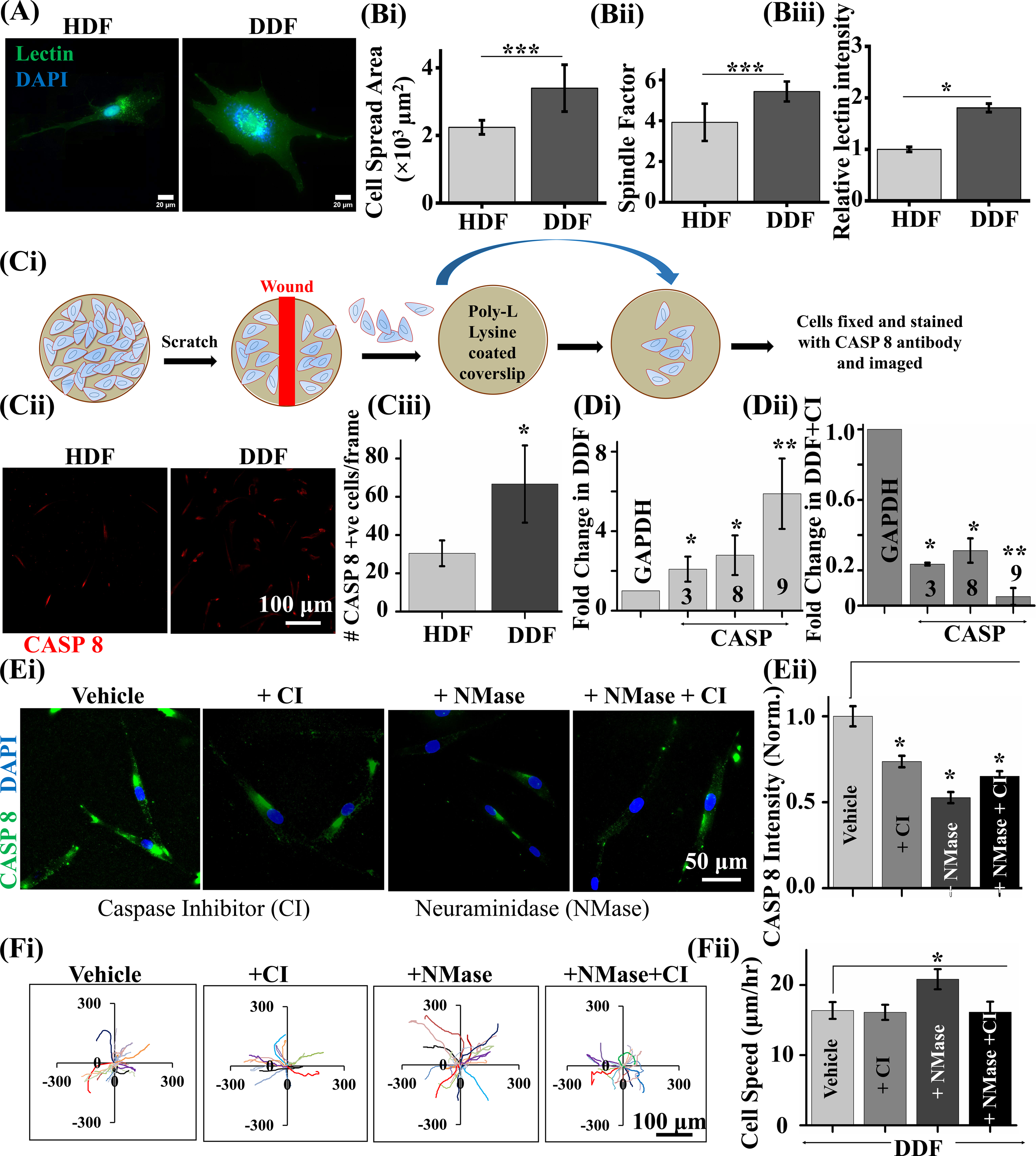
Probing the apoptotic nature of dermal fibroblast in diabetic conditions. (A) Fluorescent imaging of cell surface with lectin of the healthy dermal fibroblasts (HDFs) and diabetic dermal fibroblast (DDFs) isolated from patient skin tissues, (B) (i) Quantification of cell spread area of HDFs and DDFs (*N* = 3 and *n* ≥ 30, ‘*N*’ represents the number of independent experiments and ‘*n*’ represents the number of cells analyzed in each independent experiment). (B) (ii) quantification of the spindle factor (major axis of cell/ minor axis of cell) of HDFs and DDFs, (B)(iii) Quantification of lectin intensity at the cell periphery of the HDFs and DDFs. Statistical significance was determined using Student’s *t* test (\**p* ≤ 0.05, \*\*\**p* ≤ 0.001). Error bars represent standard deviation (± STD). (C) (i) Schematic of the wound assay for mimicking cell damage in wounds. (C) (ii, iii) representative fluorescence image of HDFs and DDFs stained with Caspase 8 and quantification of the percentage of caspase 8 positive cells (*n* ≥ 30 cells per condition, *N* = 3 independent samples). (D) (i) Quantification of Caspase 3, 8 and 9 in DDF with respect to HDF. (C) (ii) Quantification of fold change of caspase 3, 8 and 9 in DDFs after treated with Caspase inhibitor (CI) (*N* = 4 independent samples). (E) (i) Representative fluorescence images of DDFs stained with caspase 8 of the cells treated with neuraminidase (NMase) and Caspase inhibitor (CI) (*N* = 3 independent samples). (E) (ii) quantification of the fluorescent signal intensity of Caspase-8 in DDFs and DDFs treated with NMase or CI (*n* ≥ 30 cells per condition, *N* = 3). Statistical significance was determined using one-way ANOVA and the means were compared using the Tukey test (\*\*\**p* ≤ 0.001, \*\**p* ≤ 0.01, \**p* ≤ 0.05). Error bars represent standard deviation (± STD). (F)(ii) Quantification of cell speed of DDF treated with NMase or CI (*n* ≥ 30 cells per condition, *N* = 5). Error bars represent standard deviation (± SEM)

Consistent with our earlier work where we showed that DDFs possess a bulkier glycocalyx compared to HDFs [26], lectin intensity of DDFs was nearly 2-fold compared to that of HDFs (Supp. Fig. 1C). Glycocalyx disruption with neuraminidase (NMase) was shown to cause increased proliferation and increased cell contractility [26]. Interestingly, NMase was found to reduce caspase 8 (CASP8) levels similar to that of CI, with combination treatment providing no additional benefit (Fig. 1Di, ii). Furthermore, consistent with our previous study, NMase treatment led to faster motility of DDFs, with no effect with CI treatment (Fig. 1Ei, ii). Collectively, these results identify glycocalyx disruption as a novel strategy to reverse fibroblast apoptosis and stimulate their migration.

### 3.2 Chitosan-gelatin hybrid hydrogels provide better mechanical strength and antibacterial activity

Based on the above findings, we hypothesize that NMase encapsulated tissue adhesives may effective for treatment of diabetic wounds. While methacrylated gelatin (GelMA) gels have been widely used as tissue adhesives [30] [36] [33] [34], 10-15% GelMA concentrations that are widely used leads to high adhesivity with the gels also being soft [Iffat Jahan E. G., 2019]. Since diabetic wounds are prone to infection, we hypothesized that incorporation of chitosan into GelMA gels may improve mechanical properties of the gels, provide optimal adhesivity and confer protection against bacterial infections. Therefore, hybrid gels were fabricated by combining 15% GelMA with 1% low molecular weight (LMW) and high molecular weight (HMW) methacrylated chitosan (ChMA), respectively, and polymerized by subjecting the precursor gel solutions for 1 min of 365 nm UV radiation (Fig. 2Ai, ii). Both GelMA gels and the hybrid gels were hydrophilic in nature (Fig. 2C). Incorporation of ChMA led to substantial reduction in pore size (Fig. 2Aii, iii) and 8-12 fold increase in storage modulus (IJ ) of the hybrid gels (Fig. 2D). In line with greater strength, degradation kinetics revealed significantly slower degradation of the hybrid gels, with HMWH gels being the most sturdy (Fig. 2E). Furthermore, comparable anti-bacterial activity against both gram positive and gram negative bacteria were observed with both LMWH and HMWH hydrogels (Fig. 2F). When NMase was entrapped in the gels (i.e., N-GelMA, N-LMWH and N-HMWH gels), consistent with smallest pore size and high strength, slowest release of NMase was observed in HMWH gels (Fig. 2G). Together, these results suggest incorporation of ChMA enhances physical stability of hybrid gels while slowing the release kinetics of NMase, and confers antibacterial property to the hybrid gels.

**Figure 2.**
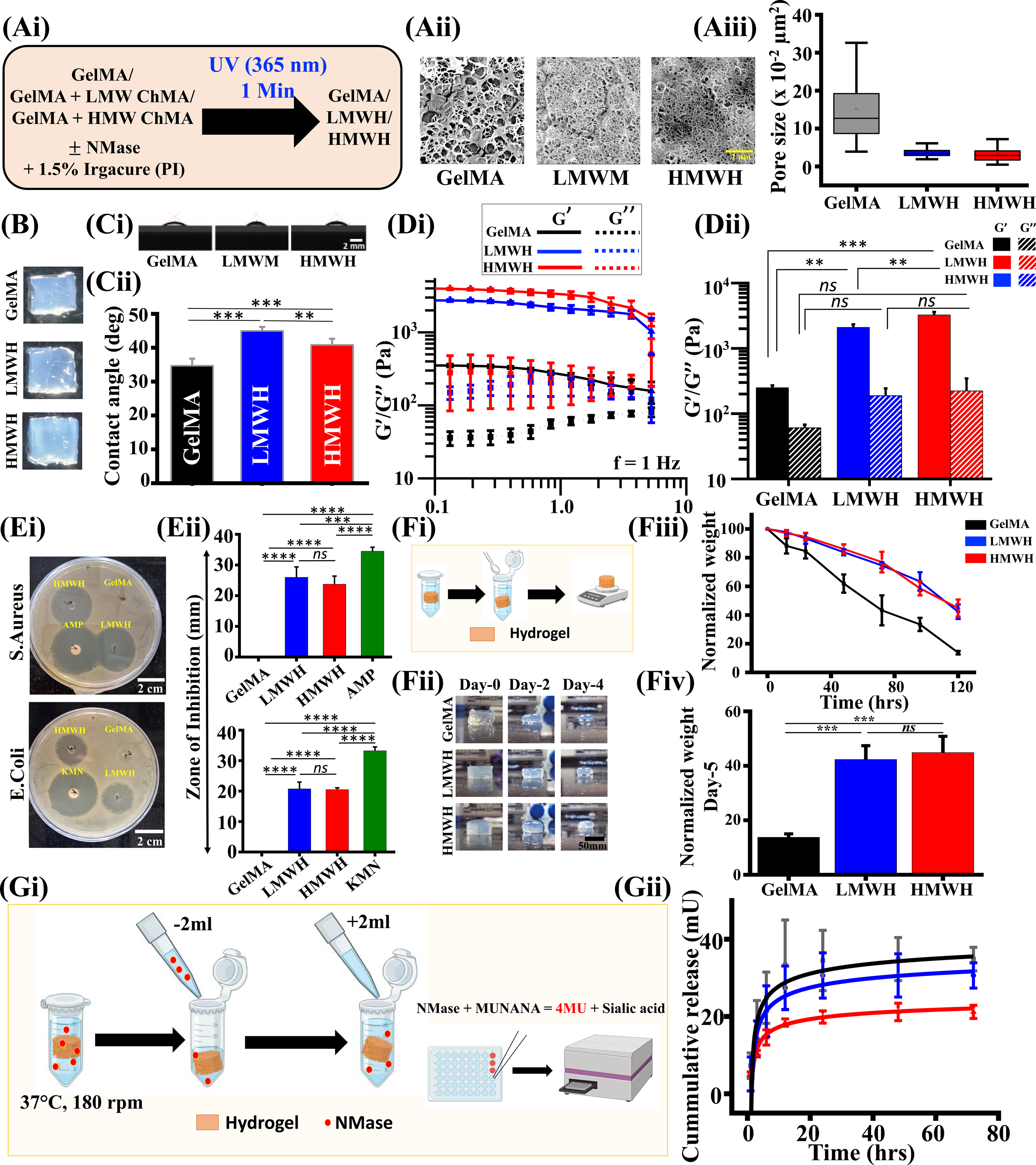
Synthesis and characterization of chitosan/gelatin hydrogels.(A)(i) Schematic of synthesis of hybrid hydrogels, (A)(ii) Cryo-FEG-SEM images of the cross-section of the hydrogels, (A)(iii) Quantification of the pore size of the hydrogels (*n* = 50 pores, *N* = 2, independently prepared gels), (B) Digital images of the hydrogel taken using CDOS camera (C)(i) Digital images of gel precursor droplet taken at 90° angle, (C)(ii) Quantification od contact images of the gel precursors to measure the hydrophilicity of the gels (*N* = 5), (D)(i, ii) Amplitude sweep of the hydrogels performed using parallel plate rheometer showing the linear viscoelastic range of the hydrogels at a frequency of 1Hz (*N* = 3 independent samples per condition) and quantification of the storage and loss moduli of the hydrogels at 1% strain, (E)(i) Representative images of disk diffusion assay taken after 16 hours of incubation with S. Aureus and E.coli bacterial strains, (E)(ii) Quantification of the zone of inhibition for the gels and control diffusion disks for S. Aureus and E.coli bacterial strains. Ampicilin (AMP) and kanamycin (KN) served as positive controls, (F)(i-iv) Schematic of enzymatic degradation assay, representative images of the hydrogel buttons taken at regular time intervals, and quantification of degradation kinetics (*N=3* independent samples per condition and statistical significant was done using Student’s *t* test (*** *p* ≤ 0.001)), (G) (I, ii) Schematic of NMase release kinetics experiment with MUNANA assay, and quantification of cumulative NMase release from the hydrogels (*N* = 3 independent samples per condition and error bars represent standard error of mean (±SEM)). Statistical significance was determined using one-way ANOVA and the means were compared using the Tukey test (\*\*\*\**p* ≤ 0.0001, \*\*\**p* ≤ 0.001, \*\**p* ≤ 0.01, \**p* ≤ 0.05). Error bars represent standard deviation (± STD).

### 3.3 NMase loaded gels enhance motility and proliferation of DDFs

We have previously demonstrated that NMase-treated DDFs exhibit enhanced motility through the regulation of focal adhesions and increased cellular contractility [26]. To evaluate the efficacy of NMase-loaded gels in boosting cell motility and proliferation, DDFs were seeded on GelMA and hybrid gels fabricated in the presence and absence of NMase. While cell viability remained unaltered on both GelMA and hybrid gels with and without NMase (Fig. S3Ai, ii), cell proliferation assessed using Ki67 staining revealed increased proliferation in the presence of NMase on both GelMA and hybrid gels, with highest proliferation rate observed on HMWH gels (Fig. 3Ai, ii).

**Figure 3.**
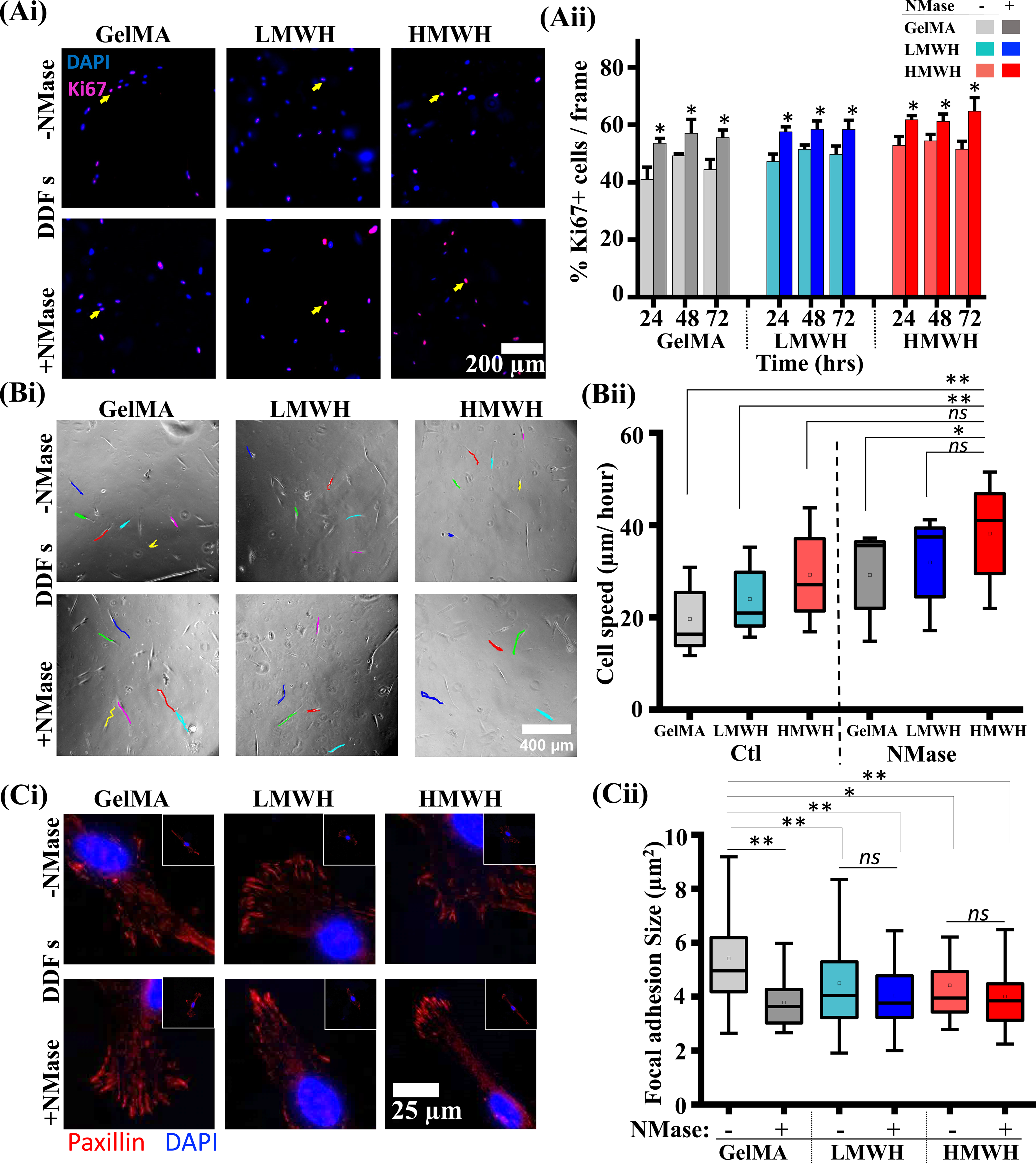
Proliferation, motility and Focal adhesion size quantification of DDFs seeded on GelMA, LMWH and HWWH gels +/- NMase. (A) (i, ii) Representative fluorescence images of DDFs stained for Ki67 protein (magenta) and nuclei (blue) and quantification of percentage Ki67 positive cells per frame (*N* = 3 from cells isolated from individual diabetic patients), (B) (I, ii) Representation single cell trajectories obtained from the time lapse images of the cells in phase contrast and quantification of cell speed (*N* = 3 and *n* ≥35)., (C) (I, ii) Representative fluorescent images of DDFs stained with paxillin antibody to measure focal adhesions and quantification of focal adhesion size of DDFs seeded on hydrogels (*N* = 3 and *n* ≥30). Statistical significance was determined using one-way ANOVA and the means were compared using the Tukey test (\*\**p* ≤ 0.01, \**p* ≤ 0.05) and the error bars represent the standard deviation (± STD).

To evaluate cell migration, time-lapse imaging was performed for 12 hours to track the movement of individual cells (Fig. 3Bi). While there was considerable variation in cell speed in each group, we saw a consistent increase in cell speed on LMWH and HMWH gels. This trend was also observed on NMase loaded gels with cells cultured on N-HWMH gels exhibiting faster migration compared to those on N-LMWH and N-GelMA gels (Fig. 3Bii). We further stained the cells for focal adhesion (FA) using paxillin antibody by fixing the cells at 24 hours after seeded on the hydrogels. We saw a decreases in the average focal adhesion (FA) size in the cells seeded on the NMase gels. There was significant decrease in FA size on the N-GelMA and N-LMWH (Fig 3Ci, ii) compared to the blank gels while the N-HMWH gels did not show a significant decrease. Together, these results suggest that both incorporation on chitosan and incorporation of NMase promotes cell proliferation and migration with maximum combinatorial effect observed on N-HMWH gels.

### 3.4 N-HMWH gels drive efficient wound healing in vivo

To test our in-vitro results in vivo, wound healing experiments were performed in a diabetic rat model. Diabetes was induced in Sprague-Dawley rats on a high fat diet for 8 weeks and by administering a single dose of Streptozotocin (STZ) after the 4^th^ week (Fig. 4Ai). Upon diabetes induction, the animals were prepped for the wound healing experiments after being anesthetized with ketamine and xylazine. The wound was created by making 1x 1 cm excision on the dorsal region. Lyophilized 1 x 1 cm hydrogel patches were hydrated in PBS for 10 minutes before being applied on the wounds. After placing the gels on the wound, Tegaderm breathable films were placed over the wound area to make sure the gels did not get detached or were removed by the animals upon regaining consciousness. The films were removed after 3-4 days once the animals adjusted to the wounds. Experiments were performed with 6 groups: control or no treatment (Group 1), GelMA and HMWH gels without NMase (Groups 2, 3) N-GelMA and N-

**Figure 4.**
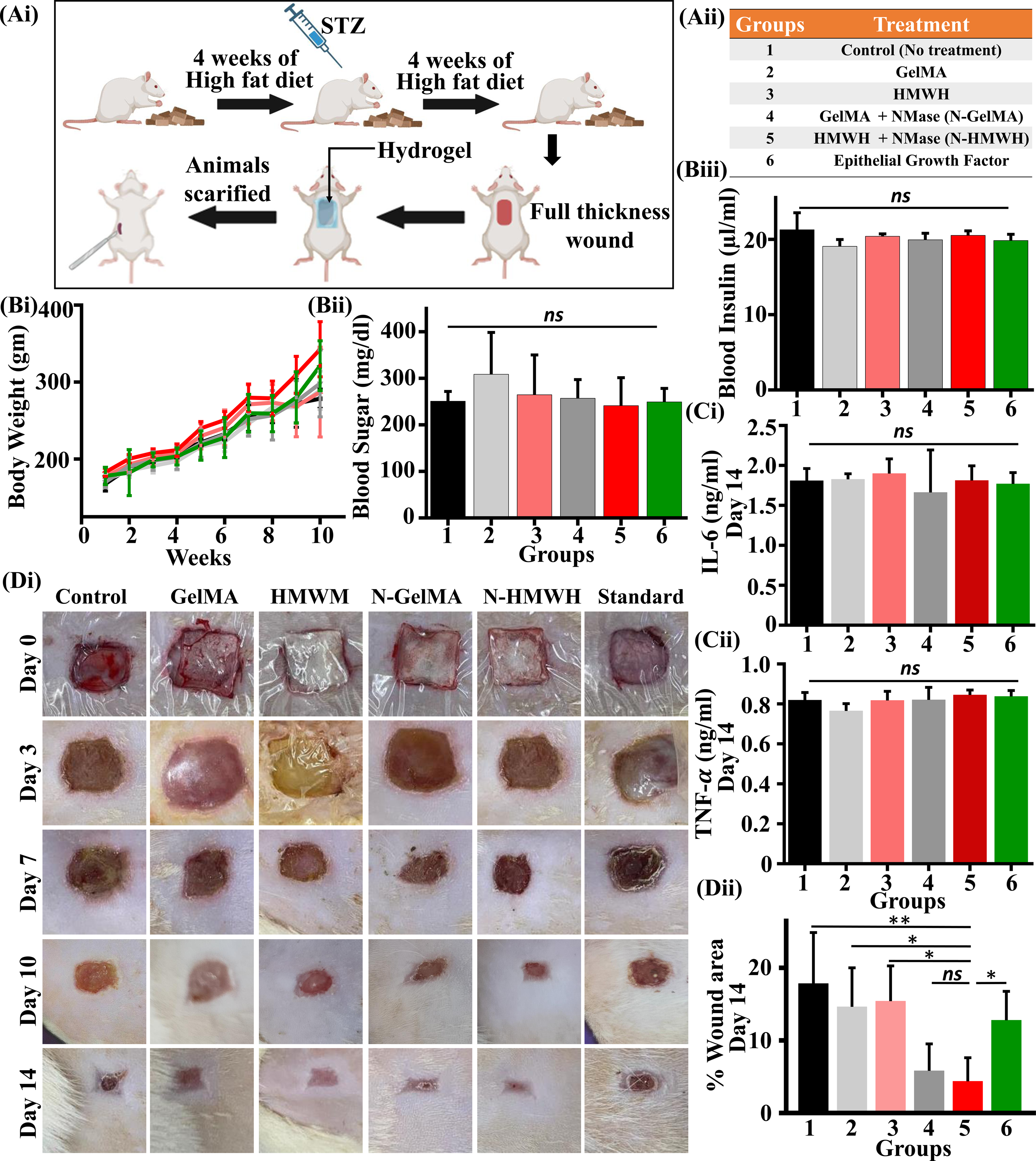
In vivo efficacy of GelMA, LMWH and HWWH gels +/- NMase gels using diabetic rat model. (A)(i) Schematic of the induction of diabetes in Sprague-dawley rats. (A)(ii) list of study groups used for wound healing experiments, (B)(i) Quantification of the body weight of the rats through the duration of the diabetic induction where the rats were under high fat diet (*N* = 6 animals per group), (B)(ii) The blood sugar levels of the rats one month after the administering of the STZ dosage (*N* = 5 animal per group), (B)(iii) blood insulin level of the rats after one month of administering STZ dosage (*N* = 5 animals per group), (C)(i) IL-6 level in the plasma of the rats on day-14 of the wound healing experiment (*N* = 5 animals per group), (C)(ii) TNF-α level in the plasma of the rats on day-14 of the wound healing experiment (*N* = 5 animals per group), (D)(i) Digital images of the wound area of all the groups taken at regular intervals, (D)(ii) Quantification of the relative wound area with respect to the wound area at day 0 (*N* = 5 animals per group). Statistical significance was determined using one-way ANOVA and the means were compared using the Tukey test (\*\**p* ≤ 0.01, \**p* ≤ 0.05). Error bars represent standard deviation (± STD).

HWMH gels which contained NMase (Group 4, 5), and animals treated with Epidermal growth factor (EGF) (Group 6) (Fig. 4Aii). Irritation studies performed using the CAM assay revealed no visible irritation to the blood vessels (Supp. Fig S4). Body weight, blood sugar levels and inflammatory markers remained unchanged across the conditions (Fig. 4B, C). Remarkably, at Day-14, the wounds were closed to the greatest extent in animals treated with NMase entrapped gels, i.e. N-GelMA and N-HMWH (Fig. 4D).

To obtain insights into why NMase loaded gels led to faster wound closure, H & E staining was performed at Day 14 to assess the formation of the epidermis layer and the formation of new blood vessels (magenta). Fastest wound healing observed in animals treated with N-HMWH gels revealed highest epidermal thickness and blood vessel density in these animals (Fig 5A). Tissue sections were also treated with Masson trichrome to assess collagen deposition. In the Masson trichrome stain, we were able to see newly formed skin appendages. There was increased collagen secretion in all the treatment conditions compared to the control (no treatment). The N-HMWH and HMWH conditions showed higher collagen fraction compared to the N-GelMA and GelMA conditions, respectively (Fig 5B). Collectively, our results suggest that faster diabetic wound healing was associated with increased collagen deposition, more complete epidermal regeneration and increased vasculature.

**Figure 5.**
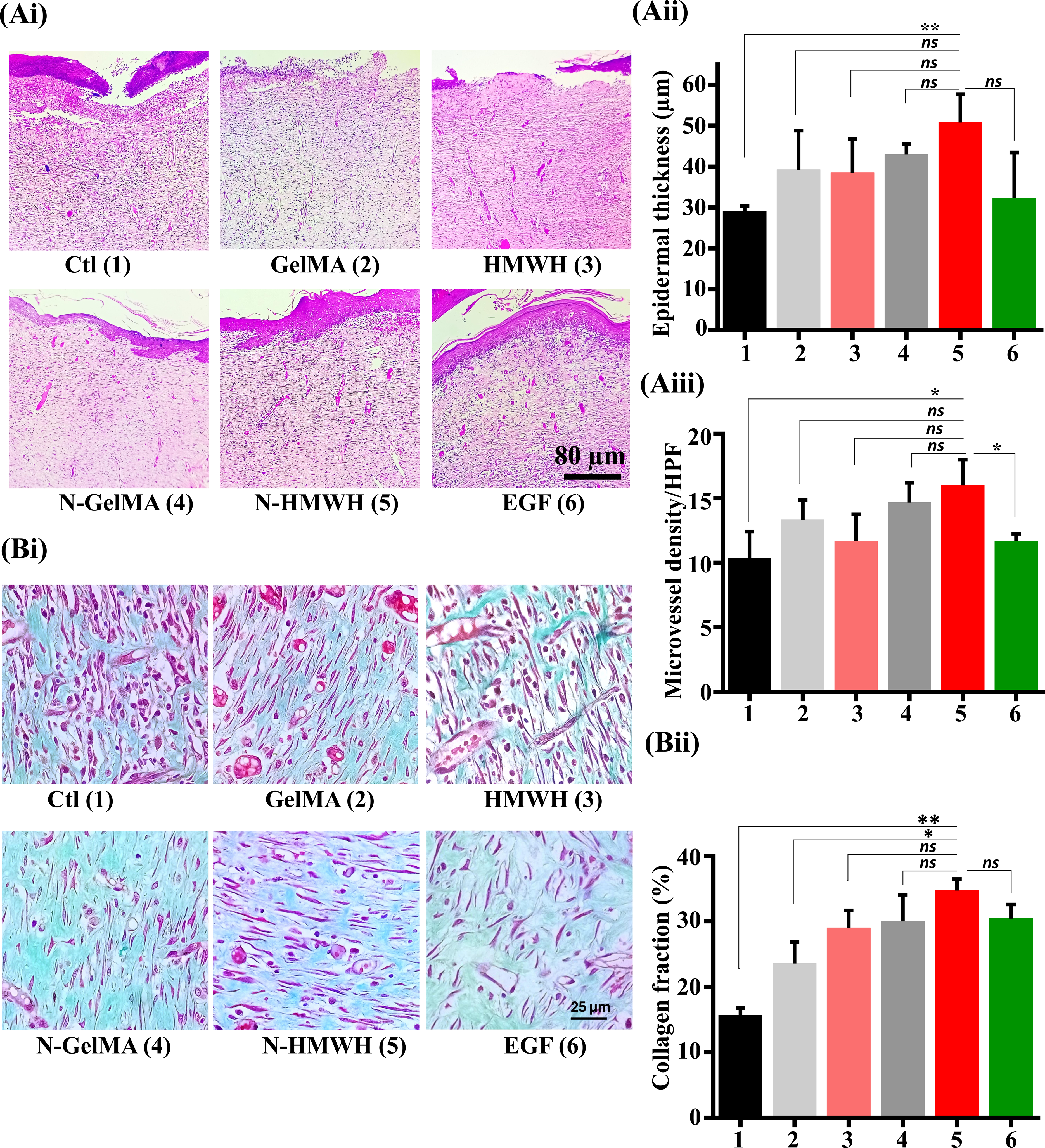
(A)(i) Hematoxylin and Eosin staining of wound tissues excised at Day 14. Purple color represents cell nuclei and the pink color represents the cytosol of the cells and the extracellular matrices. (A)(ii) The quantification of the epidermis thickness of the groups from the H&E staining, Re-epithelialization is a metric that can be used to quantify the regeneration during wound healing (*N* = 3 samples per condition), (A)(iii) Quantification of the microvessel density from the H&E staining (*N* = 3 samples per condition), (B)(i) Masson Trichrome staining of the wound granulation tissue on day-14, (B)(ii) Quantification of the collagen deposit from the Masson trichrome images where the blue color represent the ECM, the collagen deposition is measure from the intensity of the blue color from the images (*N* = 3 samples per condition). Statistical significance was determined using one-way ANOVA and the means were compared using the Tukey test (\*\**p* ≤ 0.01, \**p* ≤ 0.05). Error bars represent standard deviation (± STD).

## 4. Discussion

Wound healing depends on the coordinated progression of angiogenesis, extracellular matrix deposition and contraction to restore tissue architecture and function [9] [10] [11]. In diabetic wounds, these processes are impaired due to compromised fibroblast activity, insufficient vascularization, and persistent inflammation, resulting in increases fibroblast apoptosis, delayed healing and defective tissue remodeling [4] [14] [12] [37]. Here we show enhanced angiogenesis and collagen deposition in diabetic rats treated with NMase loaded GelMA–ChMA hydrogels, particularly in the HMWH group; this is likely attributed to the combined effects of improved mechanical stability, enhanced bioactivity, and the sustained release of NMase compared with GelMA alone. Incorporation of ChMA increases scaffold stiffness, slows degradation and provides protection against bacterial infections, thereby providing a stable microenvironment conducive to fibroblast migration, angiogenesis, and ECM remodeling during the proliferative phase of wound healing. Moreover, the bioactive properties of NMase combined with the porous architecture of the hybrid hydrogels likely facilitate improved drug retention, nutrient transport, and cellular infiltration, thereby promoting granulation tissue formation and neovascularization. Given the central role of fibroblasts in regulating both ECM deposition and angiogenic signaling, the amelioration of fibroblast dysfunction in diabetic conditions likely contributed to the increased collagen organization and vascularization observed in the hybrid hydrogel-treated wounds.

Fibroblasts in diabetic wounds are exposed to a pathological microenvironment characterized by persistent hyperglycemia, elevated oxidative stress, chronic inflammation, and impaired extracellular matrix (ECM) remodeling, all of which contribute to aberrant cellular behavior such as premature apoptosis or fibrosis. Increased levels of reactive oxygen species (ROS) in diabetic tissues impair fibroblast survival, proliferation, and migration, thereby disrupting the normal wound healing cascade [38] [39] [40] [41]. While glycocalyx plays a protective role against ROS and shear force in endothelial cells [42] [43], in DDFs possessing an enhanced glycocalyx layer that might not be the case. The glycocalyx, primarily composed of glycosaminoglycan, proteoglycans, and glycoproteins, plays an important role in regulating cell adhesion, mechanotransduction and receptor signaling. In several pathological systems, glycocalyx has been shown to promote cell survival, such as protecting cancer cells from damage during metastasis [44] [45] [21]. However, our observations suggest that in DDFs, excessive glycocalyx accumulation may in fact impair fibroblast functionality [26]. A previous study has reported that diabetic fibroblasts exhibit defective focal adhesion formation and reduced phosphorylation of focal adhesion kinase (FAK), particularly in response to growth factors such as transforming growth factor-β (TGF-β) and fibroblast growth factor (FGF) [46], resulting in impaired adhesion, migration, and increased apoptosis. In this context, NMase-mediated glycocalyx modulation may restore fibroblast activity by cleaving the terminal sialic acid residues, thereby exposing surface adhesion molecules and improving integrin-mediated cell–substrate interactions. This enhanced accessibility of adhesion receptors could promote focal adhesion assembly, increase FAK activation, and subsequently improve fibroblast migration and survival. Collectively, these findings suggest that although glycocalyx may exert protective functions in several cellular systems, its excessive accumulation in diabetic fibroblasts hinder the adhesive and regenerative processes necessary for efficient wound healing, highlighting glycocalyx remodeling as a potential therapeutic strategy for diabetic wound repair.

Methacrylated gelatin (GelMA) has been extensively utilized in tissue engineering applications due to its excellent biocompatibility, tunable physicochemical properties, and ability to support cellular adhesion and proliferation [47] [33] [31] [48]. However, the relatively low mechanical strength and rapid degradation rate of GelMA hydrogels limit their application in long-term tissue regeneration. To address these limitations, GelMA has frequently been combined with secondary polymers such as alginate, polyethylene glycol (PEG), gellan gum [34] and chitosan to fabricate hybrid hydrogels with enhanced mechanical and biological properties [47] [36]. In the present study, methacrylated chitosan (ChMA) was incorporated with GelMA to develop hybrid hydrogels possessing improved structural stability and antibacterial properties, which are particularly advantageous for diabetic wound healing applications where the risk of infection is significantly elevated. The hybrid hydrogels were synthesized through photo-crosslinking of methacrylated gelatin and methacrylated chitosan in the presence of a photoinitiator, resulting in the formation of a dense and interconnected polymeric network. The incorporation of ChMA not only improved the mechanical stability of the hydrogels but also facilitated enhanced NMase entrapment within the hydrogel matrix. Although the two hybrid formulations exhibited comparable performance in the in vitro studies, rheological analysis revealed that the high molecular weight hydrogel (HMWH) possessed a higher storage modulus than the low molecular weight hydrogel (LMWH), potentially due to increased polymer chain length and greater crosslinking density. In contrast, no significant difference was observed in the loss modulus between the two formulations, indicating similar viscous dissipation behavior under shear deformation. Furthermore, scanning electron microscopy (SEM) analysis demonstrated comparable microstructural morphology and porosity in both hydrogels. The similar viscoelastic dissipation observed in both formulations may be attributed to polymer chain entanglement within the HMWH network, where the longer chitosan chains contribute to mechanical reinforcement while simultaneously limiting the formation of additional effective crosslinks through steric hindrance.

NNMase-mediated glycocalyx disruption significantly improved the functional behavior of DDFs as evidenced by enhanced migration, proliferation, and increased phosphorylated myosin light chain (pMLC) expression [26]. Elevated pMLC levels indicate enhanced acto-myosin contractility, which plays a critical role in generating traction forces required for cell migration and extracellular matrix (ECM) remodeling during wound healing. Previous studies have reported that diabetic fibroblasts possess an abnormally thick glycocalyx layer that stabilizes focal adhesions and restricts efficient cell motility [26]. The decrease in the size of focal adhesion could mean more nascent focal adhesions form. NMase-induced desialylation likely promotes focal adhesion turnover and improves integrin-mediated mechanotransduction thereby facilitating faster cellular migration. In addition, improved proliferation following NMase treatment suggests restoration of fibroblast activity that is typically impaired under diabetic conditions. Since these effects were consistently observed across different hydrogel matrices, the findings highlight the importance of glycocalyx-mediated regulation of fibroblast mechanics and suggest that targeted glycocalyx modulation may serve as a promising therapeutic strategy for enhancing diabetic wound healing.

The role of NMase in tissue regeneration and angiogenesis remains highly context-dependent, with previous studies reporting both pro-angiogenic and anti-angiogenic effects. Angiogenesis is a tightly regulated process governed by the coordinated activity of soluble growth factors such as vascular endothelial growth factor (VEGF), fibroblast growth factor (FGF), angiopoietin-1, and thrombospondin, which collectively regulate endothelial cell proliferation, migration, and vascular maturation [49]; [50]; [51]. In the present study, NMase incorporation significantly enhanced fibroblast proliferation, which may indirectly contribute to improved angiogenesis and ECM remodeling. Fibroblasts are known to actively participate in the wound healing cascade through secretion of collagen and pro-angiogenic cytokines that support endothelial cell recruitment and capillary formation [6] [52] [11]. The enhanced fibroblast activity observed following NMase treatment may therefore create a more favorable microenvironment for tissue regeneration by promoting both matrix deposition and paracrine signaling associated with neovascularization. Furthermore, increased collagen secretion and ECM remodeling may provide structural support for newly forming vasculature within the hydrogel constructs. Collectively, these findings suggest that although the effects of NMase on angiogenesis may vary depending on the biological context, controlled glycocalyx modulation within the hydrogel system appears to promote fibroblast-mediated regenerative responses that are beneficial for diabetic wound healing.

In conclusion, our findings establish NMase-loaded hybrid hydrogels as a potent therapeutic platform for diabetic wound healing. By simultaneously modulating fibroblast glycocalyx architecture and providing mechanically optimized scaffolds (Fig. 6), this approach addresses both biochemical and biophysical deficiencies inherent to the diabetic wound microenvironment, offering a promising strategy for improved clinical outcomes.

## Acknowledgements

We acknowledge IIT Bombay for providing rheology, Cryo-FEG SEM, NMR, FTIR and Confocal Microscopy facilities. We acknowledge DST FIST supported FACS facility at IIT Bombay.

## Author Contributions

Conceptualization: VG, IJ, SSen; Methodology: VG, IJ, SD, AH, JP, RM, DK, SSen; Formal analysis: VG, IJ, AK, SR, LKB; Investigation: VG, IJ,SR, LKB, SSen.; Data curation: VG, IJ; Writing - original draft: VG, IJ, SSen; Writing - review & editing: SSen; Supervision: SSen; Project administration: SSen; Funding acquisition: SSen

