## Supplementary material for "Glycocalyx-Directed Enzymatic Hydrogels Unlocks Fibroblast Regeneration to Promote Diabetic Wound Healing": Suppl file

**Supplementary Information**

**Remodeling the Fibroblast Glycocalyx via Mechanically Tunable Hybrid Hydrogels to Accelerate Diabetic Wound Repair**

**Supp. Figure Legends**

**Supplementary Figure 1**

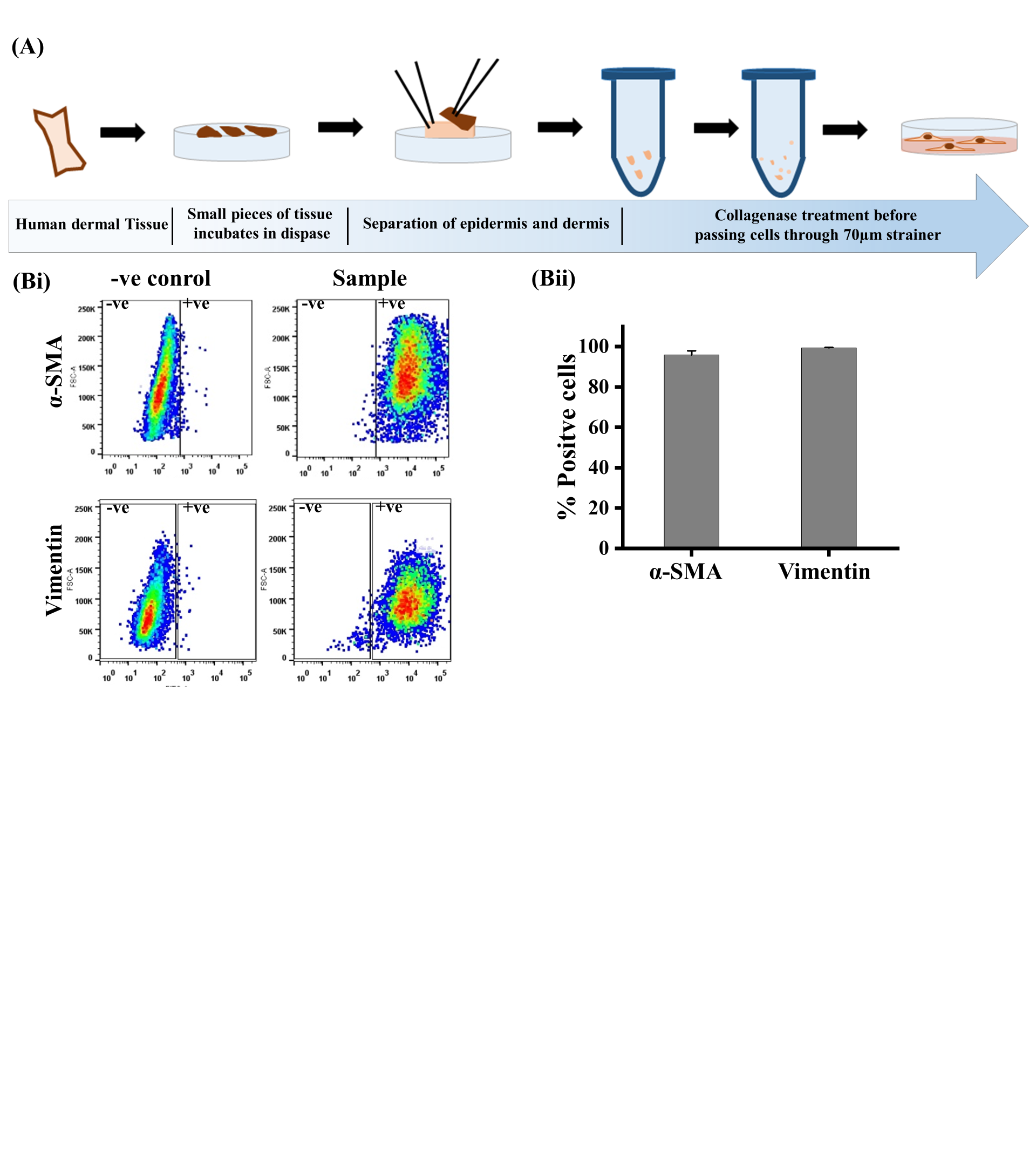

Supp. Fig. 1: Isolation and characterization of dermal fibroblasts from discarded skin tissues. (A) Schematic of the isolation of human dermal fibroblasts from skin tissue samples, (Bi) Dot plot of the isolated human dermal fibroblasts stained with α-SMA and vimentin antibodies and measured using flow cytometry, (Bii) Quantification of the percentage of cells that were α-SMA and vimentin positive, Statistical significance was determined using student t-test (*p≤0.05). Error bars represent standard deviation (± STD).

**Supplementary Figure 2:**

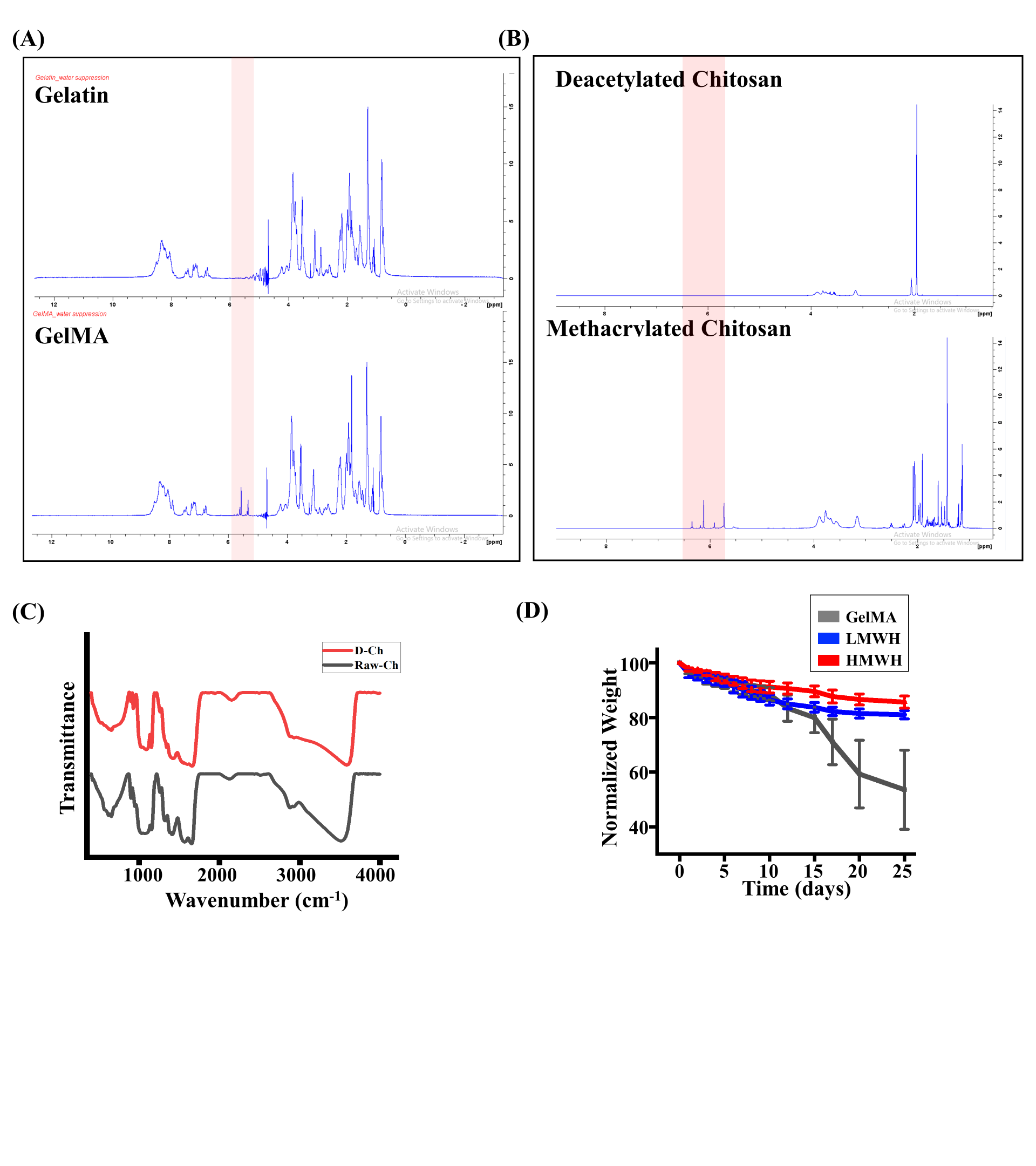

Supp Fig. 2: Characterization of chitosan, ChMA and GelMA. (A) NMR spectra of gelaitn and GelMA with highlighted peaks representing the methacrylation, with a degree of methacrytion of 65%, (B) NMR spectra of deacetylated chitosan and methacrylated chitosan (ChMA) confirming the methacrylation, with a degree of methacrytion of 35%, (C) FTIR spectrum of raw chitosan and de-acetylated chitosan with the dotted line representing the peak for the amide group, (D) Degradation kinetics of the hydrogels incubated in PBS at 37°C and shaken at 180 rpm.

**Supplementary Figure 3:**

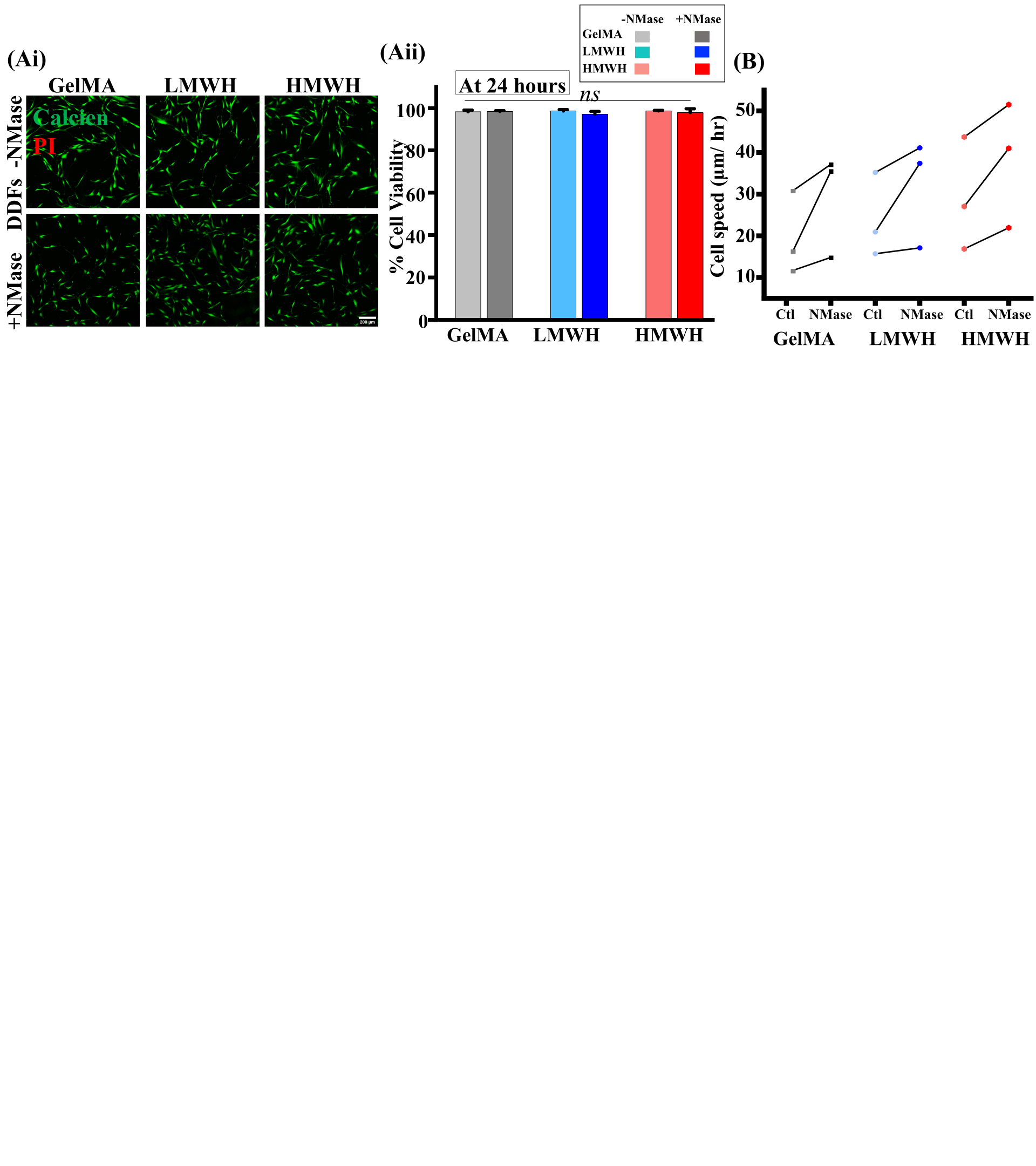

Supp Fig. 3: Cell viability and motility of DDFs on NMase hydrogels. (A i, ii) Representative fluorescence images of DDFs stained with Calcien (green) and Propidium Iodide (red) and quantification of the percentage of live cells/frame (10x magnification) (*N* = 3 from cells isolated from individual diabetic patients), (B) The connecting scatter plot representing the mean cell speed of DDFs where each dot represents the mean of single dataset Statistical significance was determined using one-way ANOVA and the means were compared using the Tukey test (**p≤0.01, *p≤0.05). Error bars represent standard deviation (± STD).

**Supplementary Figure 4:**

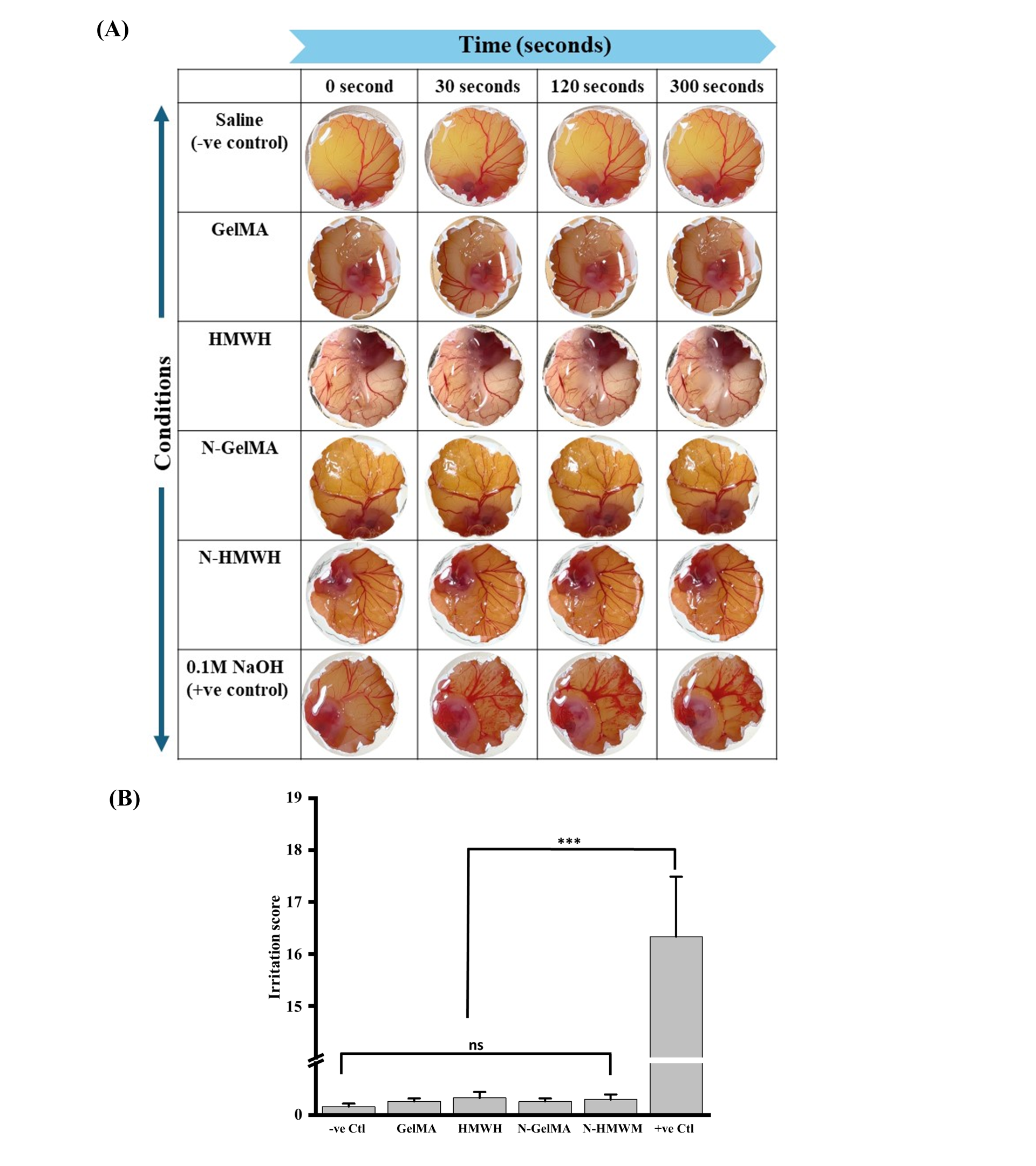

Supp Fig. 4: Irritation study of hydrogel using ex-vivo model to assess biocompatibility. (A) Representative digital images of the chick embryo chorioallantoic membrane with visible blood vessels to evaluate biocompatibility of the hydrogels, (B) Quantification of the irritation score evaluated from any haemorrhage caused by the formulations. Statistical significance was determined using one-way ANOVA and the means were compared using the Tukey test (**p≤0.01, *p≤0.05). Error bars represent standard deviation (± STD).

**Supplementary Figure 5:**

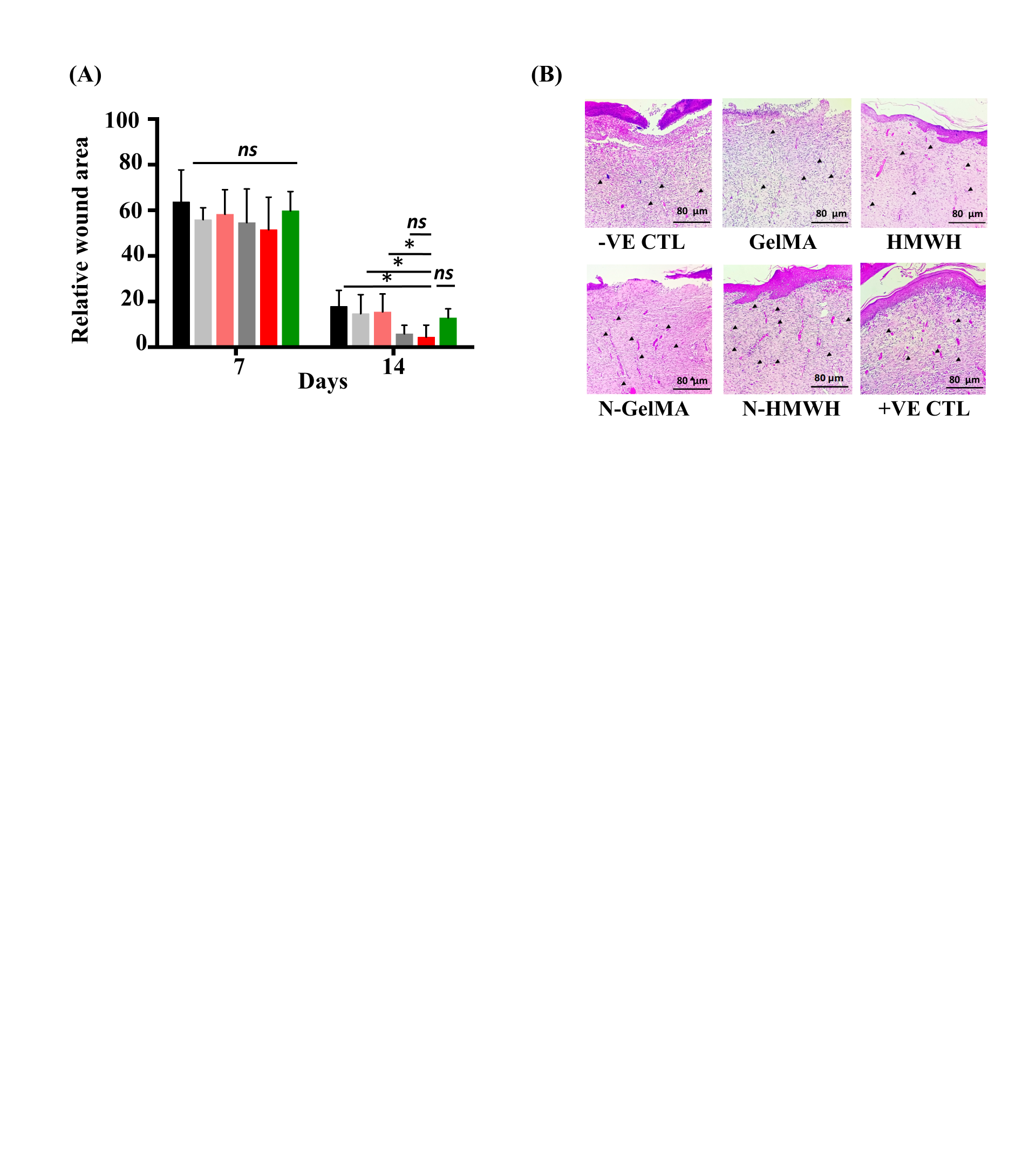
Supp Fig. 5: Quantification of wound area and H&E staining showing newly formed capillaries to assess wound healing capabilities of the NMase loaded hydrogels. (A) Quantification of the wound area on day-7 and day-14, (B) H&E staining of the excised wound tissue on day-14, where the black arrows indicate newly formed blood vessels. Statistical significance was determined using one-way ANOVA and the means were compared using the Tukey test (**p≤0.01, *p≤0.05). Error bars represent standard deviation (± STD).

**Supp. Movie Legends**

Supplementary Movie 1: Time-lapse phase contrast images of DDFs on the hydrogels to measure cell speed. Images were taken at a times of 20 minutes.
